# A Nasal Brush-based Classifier of Asthma Identified by Machine Learning Analysis of Nasal RNA Sequence Data

**DOI:** 10.1101/145771

**Authors:** Gaurav Pandey, Om P. Pandey, Angela J. Rogers, Mehmet E. Ahsen, Gabriel E. Hoffman, Benjamin A. Raby, Scott T. Weiss, Eric E. Schadt, Supinda Bunyavanich

**Author notes:** To whom correspondence should be addressed: Supinda Bunyavanich, MD, MPH, Icahn School of Medicine at Mount Sinai, 1425 Madison Avenue #1498, New York, NY 10029, USA.

## Abstract

Asthma is a common, under-diagnosed disease affecting all ages. We sought to identify a nasal brush-based classifier of mild/moderate asthma. 190 subjects with mild/moderate asthma and controls underwent nasal brushing and RNA sequencing of nasal samples. A machine learning-based pipeline identified an asthma classifier consisting of 90 genes interpreted via an L2-regularized logistic regression classification model. This classifier performed with strong predictive value and sensitivity across eight test sets, including (1) a test set of independent asthmatic and control subjects profiled by RNA sequencing (positive and negative predictive values of 1.00 and 0.96, respectively; AUC of 0.994), (2) two independent case-control cohorts of asthma profiled by microarray, and (3) five cohorts with other respiratory conditions (allergic rhinitis, upper respiratory infection, cystic fibrosis, smoking), where the classifier had a low to zero misclassification rate. Following validation in large, prospective cohorts, this classifier could be developed into a nasal biomarker of asthma.

## Introduction

Asthma is a chronic respiratory disease that affects 8.6% of children and 7.4% of adults in the United States [1]. Its true prevalence may be higher [2]. The fluctuating airflow obstruction, bronchial hyper-responsiveness, and airway inflammation that characterize mild to moderate asthma can be difficult to detect in busy, routine clinical settings [3]. In one study of US middle school children, 11% reported physician-diagnosed asthma with current symptoms, while an additional 17% reported active asthma-like symptoms without a diagnosis of asthma [2]. Undiagnosed asthma leads to missed school and work, restricted activity, emergency department visits, and hospitalizations [2, 4]. Given the high prevalence of asthma and consequences of missed diagnosis, there is high potential impact of improved biomarkers for asthma [5].

National and international guidelines recommend that the diagnosis of asthma be based on a history of typical symptoms *and objective findings* of variable expiratory airflow limitation [6, 7]. However, obtaining such objective findings can be challenging given currently available tools. Pulmonary function tests (PFTs) require equipment, expertise, and experience to execute well [8, 9]. Results are unreliable if the procedure is done with poor technique [8]. PFTs are usually not immediately available in primary care settings. Despite the published guidelines, PFTs are not done in over half of patients suspected of having asthma [8]. Induced sputum and exhaled nitric oxide have been explored as asthma biomarkers, but their implementation requires technical expertise and does not yield better clinical results than physician-guided management alone [10]. Given the above, the reality is that most asthma is still clinically diagnosed and managed based on self-report [8, 9]. This is problematic because most patients with asthma are frequently asymptomatic at the time of exam and under-perceive as well as under-report symptoms [11].

A nasal biomarker of asthma is of high interest given the accessibility of the nose and shared airway biology between the upper and lower respiratory tracts [12-15]. The easily accessible nasal passages are directly connected to the lungs and exposed to common environmental factors. Here we describe first steps toward the development of a nasal biomarker of asthma by reporting our identification of an asthma classifier using nasal gene expression data and machine learning (**Figure 1**). Specifically, we used RNA sequencing (RNAseq) to comprehensively profile gene expression from nasal brushings collected from subjects with mild to moderate asthma and controls, creating the largest nasal RNAseq data set in asthma to date. We focused on mild to moderate asthma because the waxing and waning nature of non-severe asthma render it relatively difficult to diagnose. Using a robust machine learning-based pipeline comprised of feature selection [16], classification [17], and statistical analyses [18], we identified an asthma classifier that accurately differentiates between subjects with and without mild-moderate asthma based on nasal brushings. We evaluated the classification performance of this asthma classifier on eight test sets of independent subjects with asthma and other respiratory conditions, finding that it performed with high accuracy, sensitivity, and specificity for asthma. Although this study’s focus is asthma, the pipeline described could potentially be used to develop classifiers for other phenotypes with high-dimensional data.

**Figure 1:**
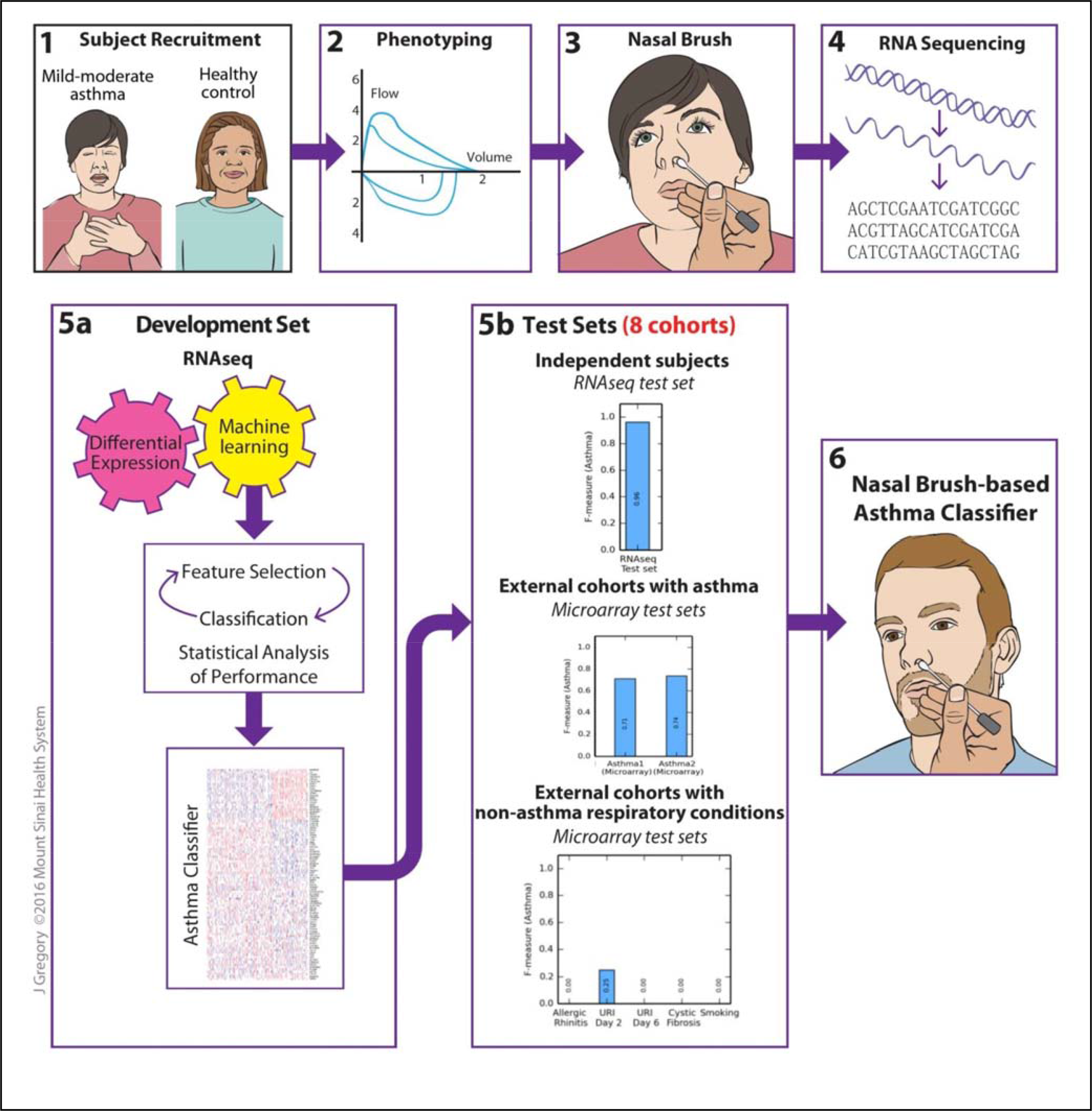
Study flow for the identification of a nasal brush-based classifier of asthma by machine learning analysis of RNAseq data. One hundred and ninety subjects with mild/moderate asthma and controls without asthma were recruited for phenotyping, nasal brushing, and RNA sequencing of nasal brushings. The RNAseq data generated were then *a priori* split into development and test sets. The development set was used for differential expression analysis and machine learning (involving feature selection, classification, and statistical analyses of classification performance) to identify an asthma classifier that can classify asthma from no asthma as accurately as possible. The asthma classifier was then evaluated on eight test sets, including (1) the RNAseq test set of independent subjects with and without asthma, (2) two external test sets of subjects with and without asthma with nasal gene expression profiled by microarray, and (3) five external test sets of subjects with non-asthma respiratory conditions (allergic rhinitis, upper respiratory infection, cystic fibrosis, and smoking) and nasal gene expression profiled by microarray. Figure drawn by Jill Gregory, Mount Sinai Health System.

## Results

### Study population and baseline characteristics

We performed nasal brushing on 190 subjects for this study, including 66 subjects with well-defined mild to moderate persistent asthma (based on symptoms, medication need, and demonstrated airway hyper-responsiveness by methacholine challenge) and 124 subjects without asthma (based on no personal or family history of asthma, normal spirometry, and no bronchodilator response). The definitional criteria we used for mild-moderate asthma are consistent with US National Heart Lung Blood Institute guidelines for the diagnosis of asthma [7], and are the same criteria used in the longest NIH-sponsored study of mild-moderate asthma [19, 20].

From these 190 subjects, a random selection of 150 subjects were *a priori* assigned as the development set (to be used for asthma classifier development), and the remaining 40 subjects were *a priori* assigned as the RNAseq test set (to be used as one of 8 test sets for evaluation of the asthma classifier identified from the development set).

The baseline characteristics of the subjects in the development set (n=150) are shown in the left section of **Table 1**. The mean age of subjects with asthma was somewhat lower than subjects without asthma, with slightly more male subjects with asthma and more female subjects without asthma. Caucasians were more prevalent in subjects without asthma, which was expected based on the inclusion criteria. Consistent with reversible airway obstruction that characterizes asthma [3], subjects with asthma had significantly greater bronchodilator response than control subjects (T-test P = 1.4 x 10^-5^). Allergic rhinitis was more prevalent in subjects with asthma (Fisher’s exact test P = 0.005), consistent with known comorbidity between allergic rhinitis and asthma [21]. Rates of smoking between subjects with and without asthma were not significantly different.

**Table 1:**
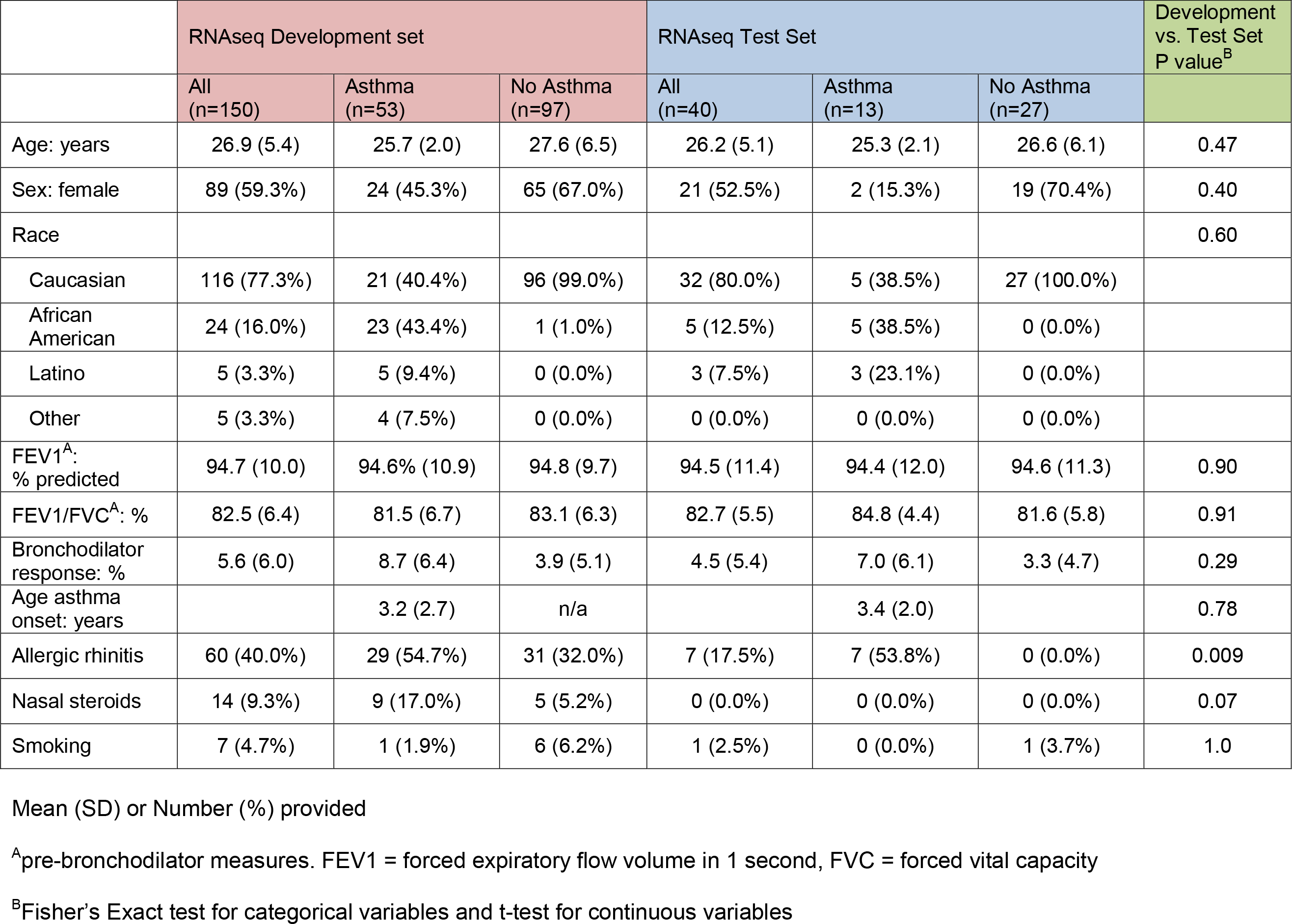
Baseline characteristics of subjects in the RNAseq development and test sets.

RNA isolated from nasal brushings from the subjects was of good quality, with mean RIN 7.8 (±1.1). The median number of paired-end reads per sample from RNA sequencing was 36.3 million. Following pre-processing (normalization and filtering) of the raw RNAseq data, 11,587 genes were used for statistical and machine learning analysis. variancePartition analysis [22], which is designed to analyze the contribution of technical and biological factors to variation in gene expression, showed that age, race, and sex contributed minimally to total gene expression variance (**Supplementary Figure 1**). For this reason, we did not adjust the pre-processed RNAseq data for these factors.

Differential gene expression analysis by DeSeq2 [23] showed that 1613 and 1259 genes were respectively over- and under-expressed in asthma cases versus controls (false discovery rate (FDR) ≤0.05) (**Supplementary Table 1**). These genes were enriched for disease-relevant pathways in the Molecular Signature Database [24], including immune system (fold change=3.6, FDR=1.07 x 10^-22^), adaptive immune system (fold change=3.91, FDR=1.46 x 10^-15^), and innate immune system (fold change=4.1, FDR=4.47 x 10^-9^) (**Supplementary Table 1**).

### Identifying a nasal brush-based asthma classifier

To identify a nasal brush-based asthma classifier using the RNAseq data generated, we developed a machine learning pipeline that combined feature (gene) selection [16] and classification techniques [17] that was applied to the development set (**Materials and Methods** and **Supplementary Figure 2**). This pipeline was designed with a systems biology-based perspective that a set of genes, even ones with marginal effects, can collectively classify phenotypes (here asthma) more accurately than individual genes [25]. More specifically, the goal of building such a classifier is not to elucidate the cause or molecular biology of the disease, but rather to identify features (genes in our study) that *in combination* can discriminate between groups of interest (e.g. asthma and no asthma). Such a classifier is likely to include genes known to associate with the groups, but it is also possible and even likely (given our incomplete understanding of complex diseases such as asthma) that genes not previously associated with the groups can provide information that is useful to the discrimination. This type of data-driven approach has been successful in other disease areas, especially cancer [26-29].

Feature selection [16] is the process of identifying a subset of features (e.g. genes) from a much larger subset in an automated data-driven fashion. In our pipeline, this process involved a cross validation-based protocol [30] using the well-established Recursive Feature Elimination (RFE) algorithm [16] combined with *L_2_*-regularized Logistic Regression (LR or Logistic) and Support Vector Machine (SVM-Linear (kernel)) algorithms [17] (combinations referred to as LR-RFE and SVM-RFE respectively) (**Supplementary Figure 3**). Classification analysis was then performed by applying four global classification algorithms (SVM-Linear, AdaBoost, Random Forest, and Logistic) [17] to the expression profiles of the gene sets identified by feature selection. To reduce the potential adverse effect of overfitting, this process (feature selection and classification) was repeated 100 times on 100 random splits of the development set into training and holdout sets. The final classifier was selected by statistically comparing the models in terms of both classification performance and parsimony, i.e., the number of genes included in the model [18] (**Supplementary Figure 4)**.

Due to the imbalance of the two classes (asthma and controls) in our cohort (consistent with imbalances in the general population for asthma and most disease states), we used F-measure as the main evaluation metric in our study [31, 32]. This class-specific measure is a conservative mean of precision (predictive value) and recall (same as sensitivity), and is described in detail in **Box 1** and **Supplementary Figure 5**. F-measure can range from 0 to 1, with higher values indicating superior classification performance. An F-measure value of 0.5 does not represent a random model. To provide context for our performance assessments, we also computed commonly used evaluation measures, including positive and negative predictive values (PPVs and NPVs) and Area Under the Receiver Operating Characteristic (ROC) Curve (AUC) scores (**Box 1** and **Supplementary Figure 5)**.

#### Box 1: Evaluation measures for classifiers

Many measures exist for evaluating the performance of classifiers. The most commonly used evaluation measures in biology and medicine are the positive and negative predictive values (PPV and NPV respectively; **Supplementary Figure 5**), and Area Under the Receiver Operating Characteristic (ROC) Curve (AUC score) [31]. However, these measures have several limitations. PPV and NPV ignore the critical dimension of sensitivity [31]. For instance, a classifier may predict perfectly for only one asthma sample in a cohort and make no predictions for all other asthma samples. This will yield a PPV of 1, but poor sensitivity, since none of the other asthma samples were identified by the classifier. ROC curves and their AUC scores do not accurately reflect performance when the number of cases and controls in a sample are imbalanced [31, 32], which is frequently the case in biomedical studies. For such situations, precision, recall, and F-measure (**Supplementary Figure 5**) are considered more meaningful performance measures for classifier evaluation [32]. Note that precision for cases (e.g. asthma) is equivalent to PPV, and precision for controls (e.g. no asthma) is equivalent to NPV (**Supplementary Figure 5**). Recall is the same as sensitivity. F-measure is the harmonic (conservative) mean of precision and recall that is computed separately for each class, and thus provides a more comprehensive and reliable assessment of model performance for cohorts with unbalanced class distributions. For the above reasons, we consider F-measure as the primary evaluation measure in our study, although we also provide PPV, NPV and AUC measures for context. Like PPV, NPV and AUC, F-measure ranges from 0 to 1, with higher values indicating superior classification performance, but a value of 0.5 for F-measure does not represent a random model and could in some cases indicate superior performance over random.

The best performing and most parsimonious combination of feature selection and classification algorithm identified by our machine learning pipeline was LR-RFE & Logistic Regression (**Supplementary Figure 4**). The classifier inferred using this combination was built on 90 predictive genes and will be henceforth referred to as the *asthma classifier*. We emphasize that the expression values of the classifier’s 90 genes must be used in combination with the Logistic classifier and the model’s optimal classification threshold (i.e. predicted label=asthma if classifier’s probability output≥0.76, else predicted label=no asthma) to be used effectively for asthma classification.

### Evaluation of the asthma classifier in an RNAseq test set of independent subjects

Our next step was to evaluate the asthma classifier in an RNAseq test set of independent subjects, for which we used the test set (n=40) of nasal RNAseq data from independent subjects. The baseline characteristics of the subjects in this test set are shown in the right section of **Table 1**. Subjects in the development and test sets were generally similar, except for a lower prevalence of allergic rhinitis among those without asthma in the test set.

The asthma classifier performed with high accuracy in the RNAseq test set’s independent subjects, achieving AUC = 0.994 (**Figure 2**), PPV = 1.00, and NPV = 0.96 (**Figures 3B and 3D, left most bar**). In terms of the F-measure metric, the classifier achieved F = 0.98 and 0.96 for classifying asthma and no asthma, respectively (**Figures 3A and 3C, left most bar**). For comparison, the much lower performance of permutation-based random models is shown in **Supplementary Figure 6**.

**Figure 2:**
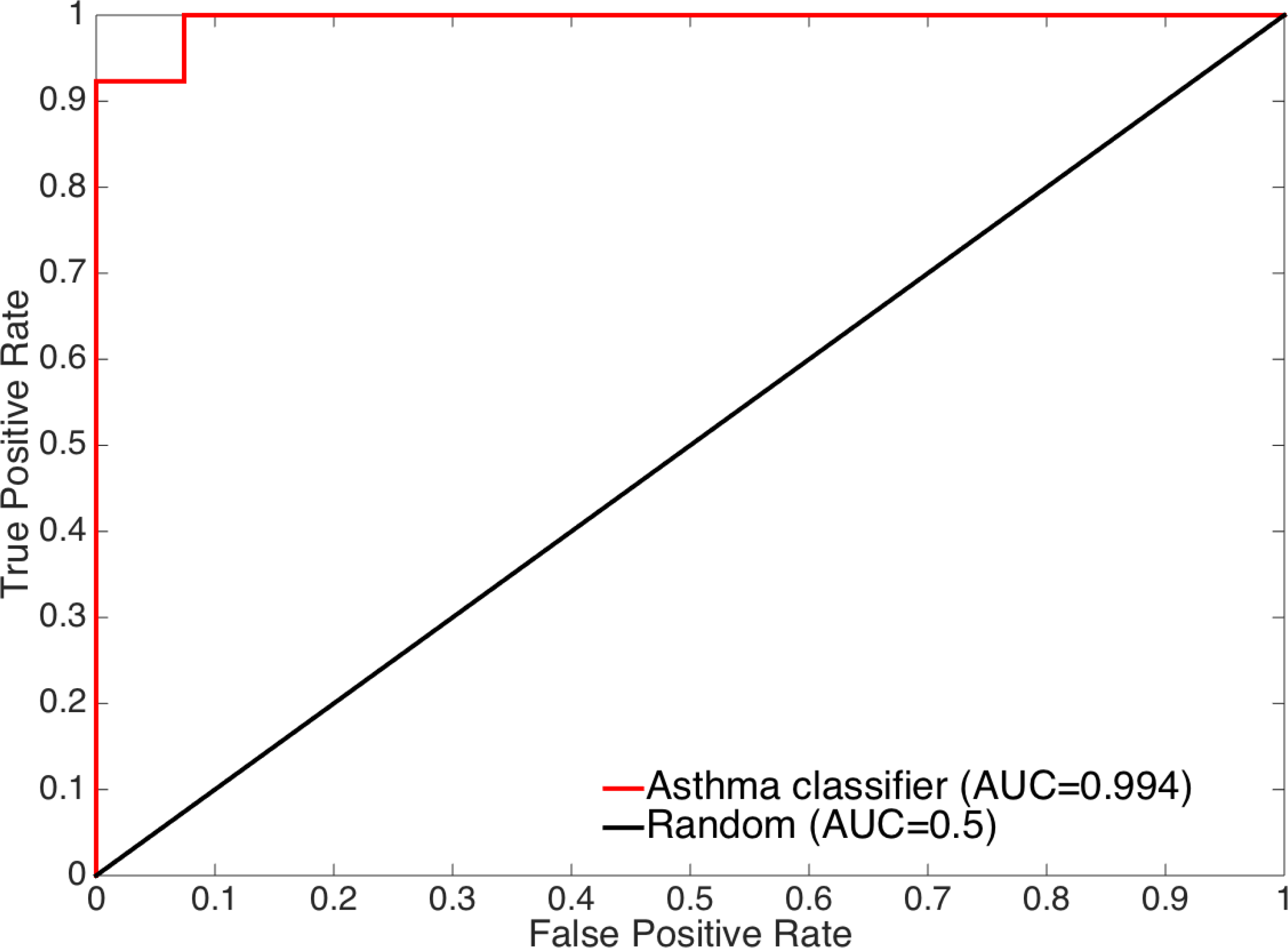
Receiver operating characteristic (ROC) curve of the predictions generated by applying the asthma classifier to the RNAseq test set of independent subjects (n=40). The ROC curve for a random model is shown for reference. The curve and its corresponding AUC score show that the classifier performs well for both asthma and no asthma (control) samples in this test set.

**Figure 3:**
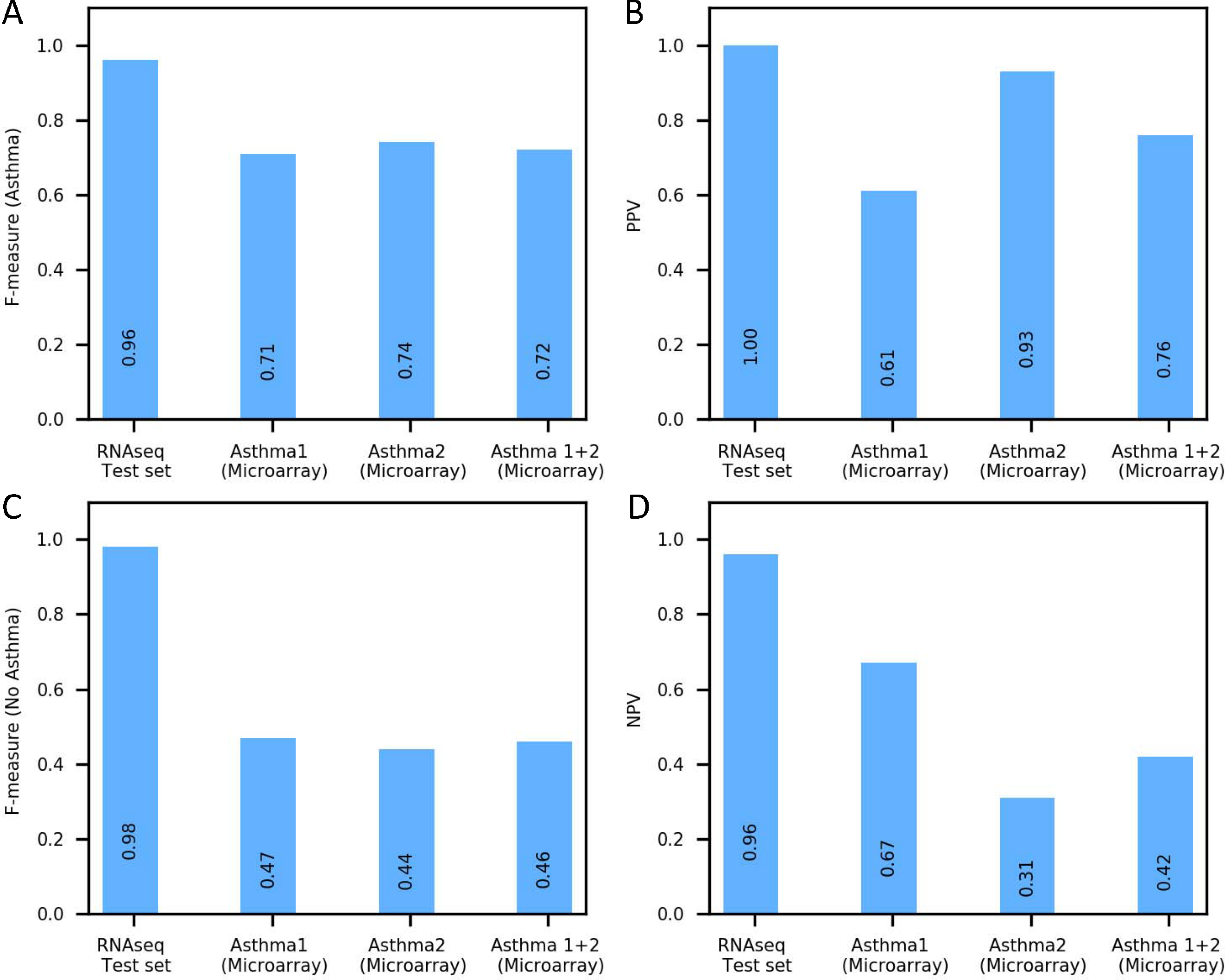
Evaluation of the asthma classifier on test sets of independent subjects with asthma. Performance of the asthma classifier in classifying asthma (A) and no asthma (C) in terms of F-measure, a conservative mean of precision and sensitivity. F-measure ranges from 0 to 1, with higher values indicating superior classification performance. The classifier was applied to an RNAseq test set of independent subjects with and without asthma, two external microarray data sets from subjects with and without asthma (Asthma1 and Asthma2), and combined data from Asthma1 and Asthma2. Positive (B) and negative (D) predictive values are also provided for context.

Our machine learning pipeline evaluated models from several combinations of feature selection and classification algorithms to select the most predictive classifier. Potentially predictive genes can also be identified from differential expression analysis and results from prior asthma-related studies. **Figure 4** shows the performance of the asthma classifier in the RNAseq test set next to alternative classifiers trained on the development set using: (1) other classifiers tested in our machine learning pipeline, (2) all genes in our data set (11587 genes after filtering), (3) all differentially expressed genes in the development set (2872 genes) (**Supplementary Table 1**), (4) genes associated with asthma from prior genetic studies [33] (70 genes) (**Supplementary Table 2**), and (5) a commonly used one-step classification model (L1-Logistic) [34] (243 genes). The asthma classifier identified by our pipeline outperformed all these alternative classifiers despite its reliance on a small number of genes.

**Figure 4:**
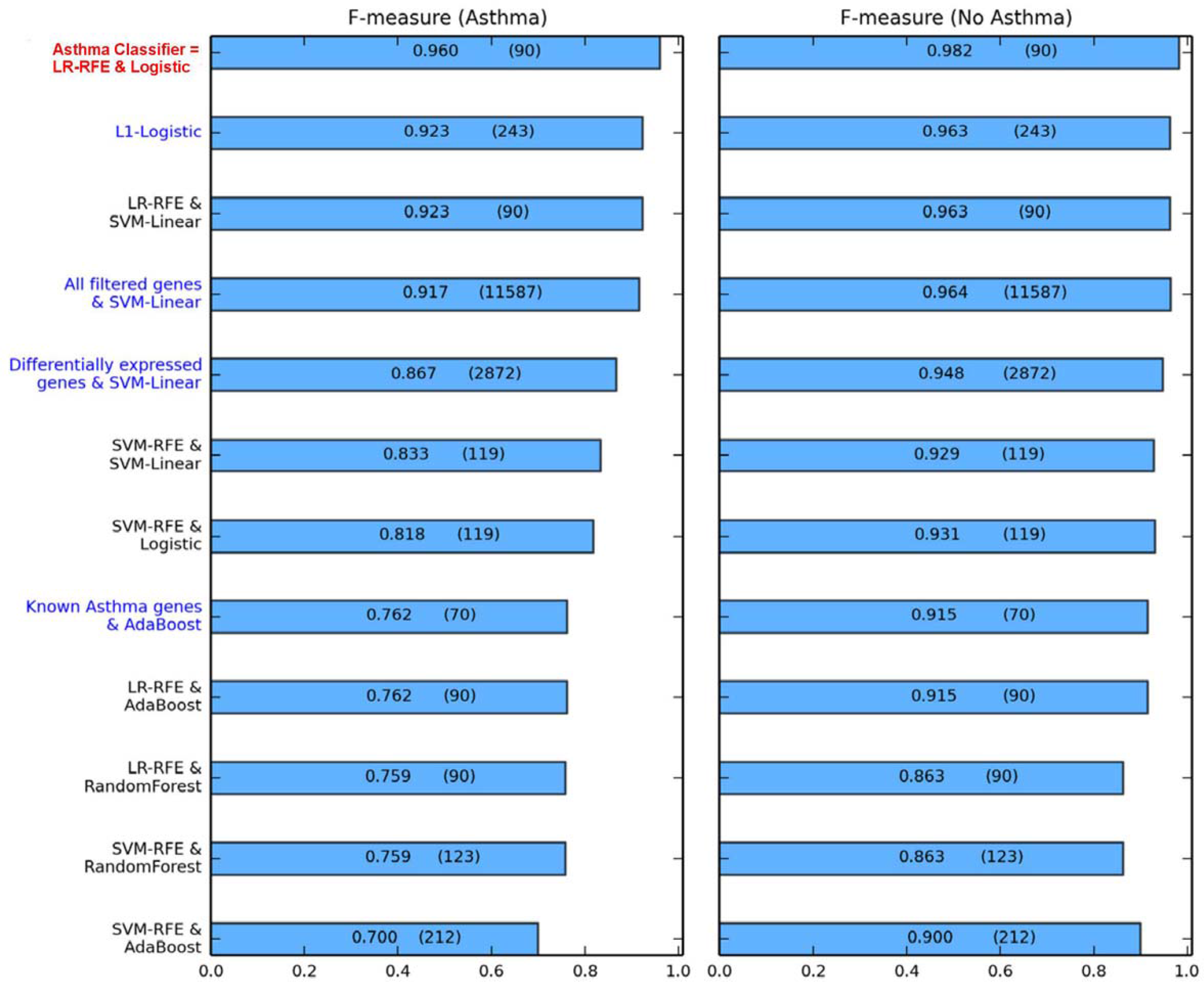
Comparative performance of the asthma classifier and other classification models in the RNAseq test set. Performances of the asthma classifier and other classification models in classifying asthma (left panel) and no asthma (right panel) are shown in terms of F-measure, with individual measures shown in the bars. The number of genes in each model is shown in parentheses within the bars. The asthma classifier is labeled in red and classification models learned from the machine learning pipeline using other combinations of feature selection and classification are labeled in black. These other classification models were combinations of two feature selection algorithms (LR-RFE and SVM-RFE) and four global classification algorithms (Logistic Regression, SVM-Linear, AdaBoost and Random Forest). For context, alternative classification models (labeled in blue) are also shown and include: (1) a model derived from an alternative, single-step classification approach (sparse classification model learned using the L1-Logistic regression algorithm), and (2) models substituting feature selection with each of 3 pre-selected gene sets (all genes after filtering, all differentially expressed genes in the development set, and known asthma genes [33]) with their respective best performing global classification algorithms. These results show the superior performance of the asthma classifier compared to all other models, in terms of classification performance and model parsimony (number of genes included). LR = Logistic Regression. SVM = Support Vector Machine. RFE = Recursive Feature Elimination.

We emphasize that our classifier produced more accurate predictions than models using all genes, all differentially expressed genes, and all known asthma genes. This supports that data-driven methods can build more effective classifiers than those built exclusively on traditional statistical methods (which do not necessarily target classification), and current domain knowledge (which may be incomplete and subject to investigation bias). Our classifier also outperformed and was more parsimonious than the model learned using the commonly used L1-Logistic method, which combined feature selection and classification into a single step. The fact that our asthma classifier performed well in an independent RNAseq test set while also outperforming alternative models lends confidence to its classification ability.

### Evaluation of the asthma classifier in external asthma cohorts

To assess the performance of our asthma classifier in other populations and profiling platforms, we applied the classifier to nasal gene expression data generated from independent cohorts of asthmatics and controls profiled by microarrays: Asthma1 (GEO GSE19187) [35] and Asthma2 (GEO GSE46171) [36]. **Supplementary Table 3** summarizes the characteristics of these external, independent case-control cohorts. In general, RNAseq-based predictive models are not expected to translate well to microarray-profiled samples [37, 38]. A major reason is that gene mappings do not perfectly correspond between RNAseq and microarray due to disparities between array annotations and RNAseq gene models [38]. Our goal was to assess the performance of our asthma classifier despite discordances in study designs, sample collections, and gene expression profiling platforms.

The asthma classifier performed relatively well (**Figure 3 middle bars**) and consistently better than permutation-based random models (**Supplementary Figure 6**) in classifying asthma and no asthma in both the Asthma1 and Asthma2 microarray-based test sets. The classifier achieved similar F-measures in the two test sets (**Figures 3A and 3C middle bars**), although the PPV and NPV measures were more dissimilar for Asthma2 (PPV 0.93, NPV 0.31) than for Asthma1 (PPV 0.61, NPV 0.67) (**Figure 3B and 3D middle bars**). The classifier’s performance was better than its random counterparts for both these test sets, although the difference in this performance was smaller for Asthma2. This occurred partially because Asthma2 includes many more asthma cases than controls (23 vs. 5), which is counter to the expected distribution in the general population. In such a skewed data set, it is possible for a random model to yield an artificially high F-measure for asthma by predicting every sample as asthmatic. We verified that this occurred with the random models tested on Asthma2.

To assess how the asthma classifier might perform in a larger external test set, we combined samples from Asthma1 and Asthma2 and performed the evaluation on this combined set. We chose this approach because no single large, external dataset of nasal gene expression in asthma exists, and combining cohorts could yield a joint test set with heterogeneity that partially reflects real-life heterogeneity of asthma. As expected, all the performance measures for this combined test set were intermediate to those for Asthma1 and Asthma2 (**Figure 3 right most bars**). These results supported that our classifier also performs reasonably well in a larger and more heterogeneous cohort.

Overall, despite the discordance of gene expression profiling platforms, study designs, and sample collection methods, our asthma classifier performed reasonably well in these external test sets, supporting a degree of generalizability of the classifier across platforms and cohorts.

### Specificity of the asthma classifier: testing in external cohorts with non-asthma respiratory conditions

To assess the specificity of our asthma classifier, we next sought to determine if it would misclassify as asthma other respiratory conditions with symptoms that overlap with asthma. To this end, we evaluated the performance of the asthma classifier on nasal gene expression data derived from case-control cohorts with allergic rhinitis (GSE43523) [39], upper respiratory infection (GSE46171) [36], cystic fibrosis (GSE40445) [40], and smoking (GSE8987) [12]. **Supplementary Table 4** details the characteristics for these external cohorts with non-asthma respiratory conditions. In three of these five non-asthma cohorts (Allergic Rhinitis, Cystic Fibrosis and Smoking), the classifier appropriately produced one-sided classifications, i.e., samples were all appropriately classified as “no asthma.” This is shown by the zero F-measure for the positive (asthma) class (**Figure 5A**) and perfect F-measure for the negative (no asthma) class (**Figure 5C**) obtained by the classifier in these cohorts. In other words, the precision for the asthma class (PPV) of our classifier was exactly and appropriately zero (**Figure 5B**), and NPV was perfectly 1.00 for these cohorts with non-asthma conditions (**Figures 5D**). The URI day 2 and 6 cohorts were slight deviations from these trends, where the classifier achieved perfect NPVs of 1.00 (**Figure 5D**), but marginally lower F-measure for the “no asthma” class (**Figure 5C**) due to slightly lower than perfect sensitivity. This may have been influenced by common inflammatory pathways underlying early viral inflammation and asthma [41]. Nonetheless, consistent with the other non-asthma test sets, the classifier’s misclassification of URI as asthma was rare and substantially less than its random counterpart classifiers (**Supplementary Figure 7**).

**Figure 5:**
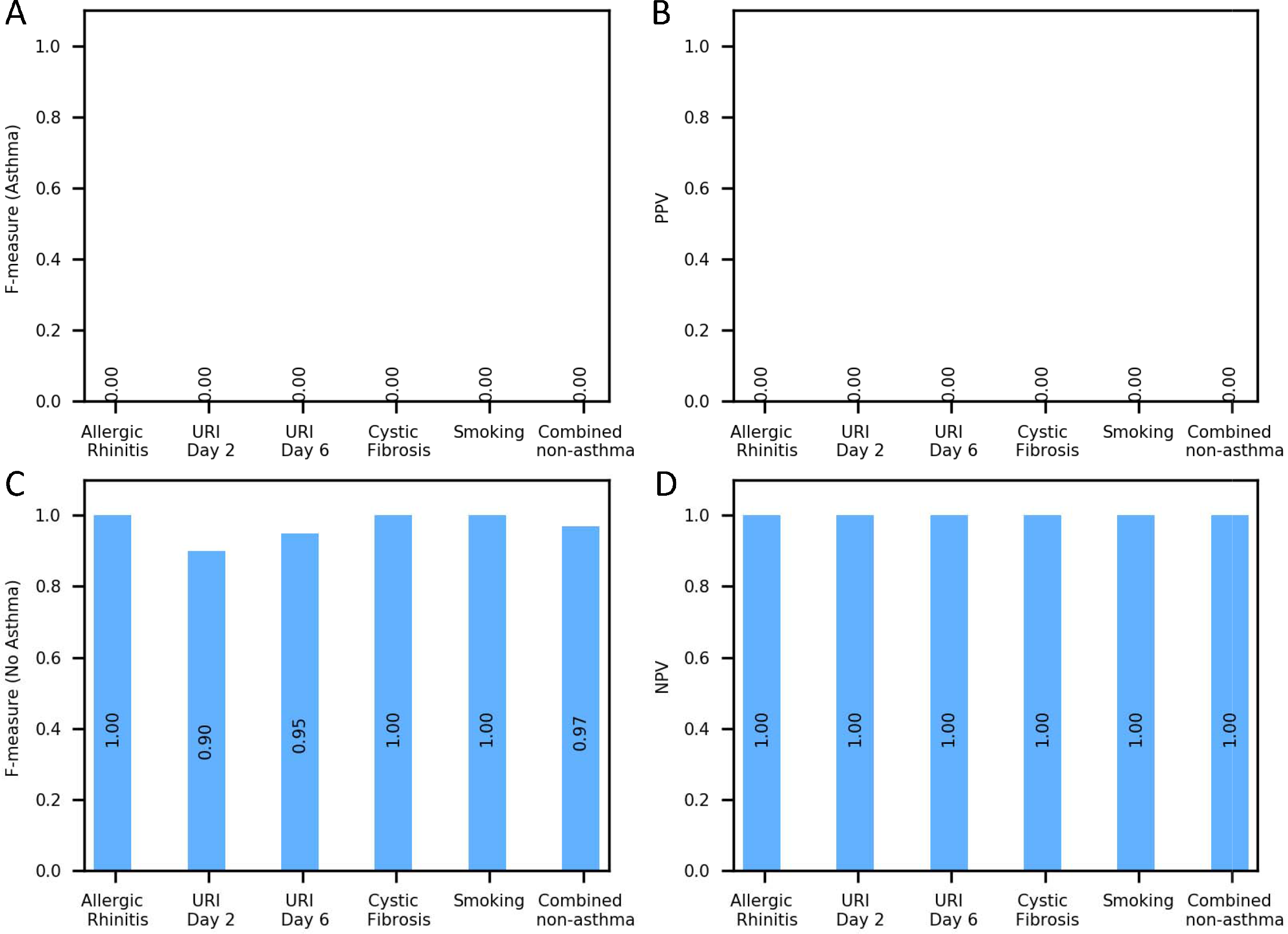
Evaluation of the asthma classifier on test sets of independent subjects with non-asthma respiratory conditions. Performance statistics of the classifier when applied to external microarray-generated data sets of nasal gene expression derived from case/control cohorts with non-asthma respiratory conditions. Performance is shown in terms of F-measure (A and C), a conservative mean of precision and sensitivity, as well as positive (B) and negative predictive values (D). The classifier had a low to zero rate of misclassifying other respiratory conditions as asthma, supporting that the classifier is specific to asthma and would not misclassify other respiratory conditions as asthma.

To assess the asthma classifier’s performance if presented with a large, heterogeneous collection of non-asthma respiratory conditions reflective of real clinical settings, we aggregated the non-asthma cohorts into a “Combined non-asthma” test set and applied the asthma classifier. The results included an appropriately zero F-measure for asthma and zero PPV, and an F-measure of 0.97 for no asthma, and NPV of 1.00 (**Figure 5, right most bars**). Results from the individual and combined non-asthma test sets collectively support that the asthma classifier would rarely misclassify other respiratory diseases as asthma.

### Statistical and Pathway Examination of Genes in the Asthma Classifier

An interesting question to ask for a disease classifier is how does its predictive ability relate to the individual differential expression status of the genes constituting the classifier? We found that 46 of the 90 genes included in our classifier were differentially expressed (FDR ≤0.05), with 22 and 24 genes over- and under-expressed in asthma respectively (**Figure 6, Supplementary Table 1**). More generally, the genes in our classifier had lower differential expression FDR values than other genes (Kolmogorov-Smirnov statistic=0.289, P-value=2.73x10^-37^) (**Supplementary Figure 8**).

**Figure 6:**
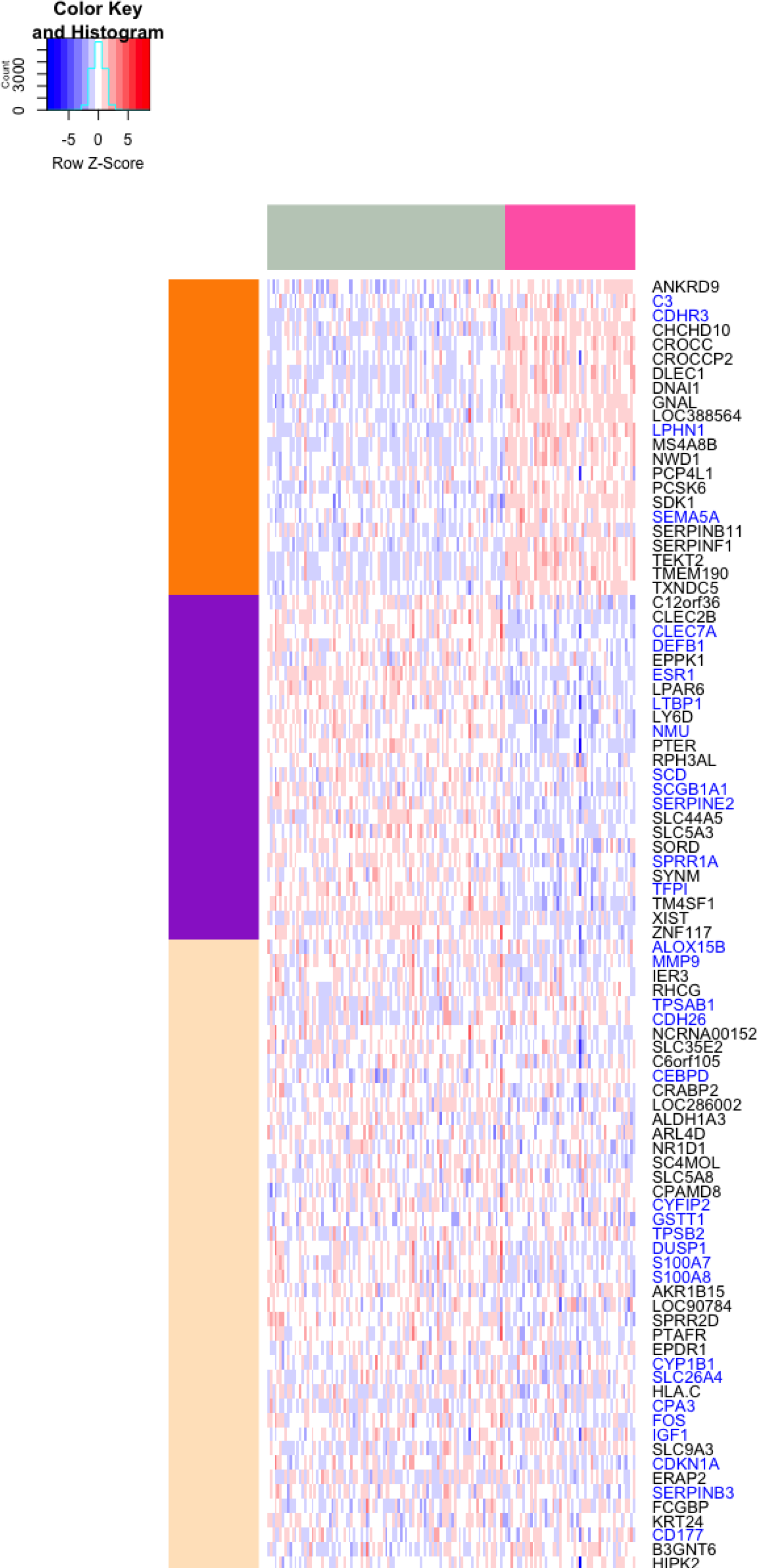
Heatmap showing expression profiles of the 90 genes constituting the asthma classifier. Columns shaded pink at the top denote asthma samples, while samples from subjects without asthma are denoted by columns shaded grey. 22 and 24 of these genes were over- and under-expressed in asthma samples (DESeq2 FDR ≤ 0.05), denoted by orange and purple groups of rows, respectively. The 33 genes in this set that have been previously studied in the context of asthma are marked in blue. The classifier’s inclusion of genes not previously known to be associated with asthma as well as genes not differentially expressed in asthma (beige group of rows) demonstrates the ability of a machine learning methodology to move beyond traditional analyses of differential expression and current domain knowledge.

In terms of biological function, pathway enrichment analysis of our classifier’s 90 genes, though statistically limited by the small number of genes, yielded enrichment for pathways including defense response (fold change=2.86, FDR=0.006) and response to external stimulus (fold change=2.50, FDR=0.012). A minority (33) of these 90 genes or their gene products have been studied in the context of asthma or airway inflammation by various modes of study as summarized in **Supplementary Table 5**. These results suggest that our machine learning pipeline was able to extract information beyond individually differentially expressed or previously known disease-related genes, allowing for the identification of a parsimonious set of genes that collectively enabled accurate disease classification.

## Discussion

Using RNAseq data generated from our cohorts, combined with a systematic machine learning analysis approach, we identified a nasal brush-based classifier that accurately distinguishes subjects with mild/moderate asthma from controls. This asthma classifier, consisting of the expression profiles of 90 genes interpreted via a logistic regression classification model, performed with high precision (PPV=1.00 and NPV=0.96) and recall for classifying asthma (AUC=0.994). The performance of the asthma classifier across independent test sets demonstrates potential for the classifier’s generalizability across study populations and two major modalities of gene expression profiling (RNAseq and microarray). Additionally, the classifier’s low to zero rate of misclassification on external cohorts with non-asthma respiratory conditions supports the specificity of this asthma classifier. Our results represent the first steps toward the development of a nasal biomarker of asthma.

Our nasal brush-based asthma classifier is based on the common biology of the upper and lower airway, a concept supported by clinical practice and previous findings [12-15]. Clinicians often rely on the united airway by screening for lower airway infections (e.g. influenza, methicillin-resistant Staphylococcus aureus) with nasal swabs [42]. Sridhar et al. found that gene expression consequences of tobacco smoking in bronchial epithelial cells were reflected in nasal epithelium [12]. Wagener et al. compared gene expression in the nasal and bronchial epithelia from 17 subjects, finding that 99% of the 33,000 genes tested exhibited no differential expression between the nasal and bronchial epithelia in those with airway disease [13]. In a study of 30 children, Guajardo et al. identified gene clusters with differential expression in nasal epithelium between subjects with exacerbated asthma vs. controls [14]. The above studies were done with small sample sizes and microarray technology. More recently, Poole et al. compared RNAseq profiles of nasal brushings from 10 asthmatic and 10 control subjects to publicly available bronchial transcriptional data, finding correlation (ρ = 0.87) between nasal and bronchial transcripts, as well as correlation (ρ=0.77) between nasal differential expression and previously observed bronchial differential expression in asthmatics [15]. To the best of our knowledge, our study has generated the largest nasal RNAseq data set in asthma to date and is the first to identify a nasal brush-based classifier of asthma.

Although based on only 90 genes, our asthma classifier classified asthma with greater accuracy than models based on all genes, all differentially expressed genes, and known asthma genes (**Figure 4**). Its superior performance supports that our machine learning pipeline successfully selected a parsimonious set of informative genes that (1) captures more actionable knowledge than traditional differential expression and genetic association analyses, and (2) cuts through the potential noise of genes irrelevant to asthma. These results illustrate that data-driven methods can build more effective classifiers than those built exclusively on current domain knowledge. This is likely true not just for asthma but for other phenotypes as well. About half the genes in our asthma classifier were not differentially expressed at FDR ≤ 0.05, and as such would not have been examined with greater interest had we only performed traditional differential expression analysis, which is the main analytic approach of virtually all studies of gene expression in asthma [12-15, 43, 44]. Consistent with motivations underlying systems biology and genomic approaches [25, 45], our study demonstrated that the asthma classifier captures signal from differential expression as well as genes below traditional significance thresholds that may still have a contributory role to asthma classification.

With prospective validation in large cohorts, our asthma classifier could lead to the development of a minimally invasive biomarker to aid asthma diagnosis at clinical frontlines, where time and resources often preclude pulmonary function testing (PFT). Nasal brushing can be performed quickly, does not require machinery for collection, and implementation of our classification model yields a straightforward, binary result of asthma or no asthma. Because it takes seconds for nasal brushing and bioinformatic interpretation could be automated, an asthma classifier such as ours may be attractive to time-strapped clinicians, particularly primary care providers at the frontlines of asthma diagnosis. Asthma is frequently diagnosed and treated in the primary care setting [46] where access to PFTs is often not immediately available. Gene expression-based diagnostic classifiers are being successfully used in other disease areas, with prominent examples including the commercially available MammaPrint [47] and Oncotype DX [48] for diagnosing breast cancer phenotypes, leading to better outcomes. These examples from the cancer field demonstrate an existing path for moving a classifier such as ours to clinical use.

We recognize that our asthma classifier did not perform quite as well in the microarray-based vs. RNAseq-based asthma test sets, which was to be expected due to differences in study design and technological factors between RNAseq and microarray profiling. First, the baseline characteristics and phenotyping of the subjects differed. Subjects in the RNAseq test set were adults who were classified as mild/moderate asthmatic or healthy using the same strict criteria as the development set, which required subjects with asthma to have an objective measure of obstructive airway disease (i.e. positive methacholine challenge response). In contrast, subjects in the Asthma1 microarray test set were all children (i.e. not adults) with nasal pathology, as entry criteria included dust mite allergic rhinitis specifically [35] (**Supplementary Table 3**). Subjects from the Asthma2 cohort were adults who were classified as having asthma or healthy based on history. As mentioned, the diagnosis of asthma based on history alone without objective lung function testing can be inaccurate [49]. The phenotypic differences between these test sets alone could explain differences in performance of our asthma classifier in these test sets. Second, the differential performance may be due to the difference in profiling approach. Gene mappings do not perfectly correspond between RNAseq and microarray due to disparities between array annotations and RNAseq gene models [38]. Compared to microarrays, RNAseq quantifies more RNA species and captures a wider range of signal [43]. Prior studies have shown that microarray-derived models can reliably predict phenotypes based on samples’ RNAseq profiles, but the converse does not often hold [38]. Despite the above limitations, our asthma classifier performed with reasonable accuracy in classifying asthma in these independent microarray-based test sets. These results support a degree of generalizability of our classifier to asthma populations that may be phenotyped or profiled differently.

An effective clinical classifier should have good positive and negative predictive value [50]. This was indeed the case with our asthma classifier, which achieved high positive and negative predictive values of 1.00 and 0.96 respectively in the RNAseq test set. We also tested our asthma classifier on independent tests sets of subjects with allergic rhinitis, upper respiratory infection, cystic fibrosis, and smoking, and showed that the classifier had a low to zero rate of misclassifying other respiratory conditions as asthma (**Figure 5**). These results were particularly notable for allergic rhinitis, a predominantly nasal condition. Although our classifier is based on nasal gene expression, and asthma and allergic rhinitis frequently co-occur [21], our classifier did not misclassify allergic rhinitis as asthma. Although these conclusions are based on relatively small test sets due to the scarcity of nasal gene expression data in the public domain, the performance of our classifier gives hope that it has the potential to be generalizable and specific in much larger cohorts as well.

Although we have generated one of the largest nasal RNAseq data set in asthma to date, a future direction of this study is to recruit additional cohorts for nasal gene expression profiling and extend validation of our findings in a prospective manner. We recognize that our development set was from a single center and its baseline characteristics, such as race, do not characterize all populations. However, we find it reassuring that the classifier performed reasonably well in multiple external data sets spanning children and adults of varied racial distributions, and with asthma and other respiratory conditions defined by heterogeneous criteria. Subjects with asthma in our development cohort were not all symptomatic at the time of sampling. The fact that the performance of our asthma classifier does not rely on symptomatic asthma is a strength, as many mild/moderate asthmatics are only sporadically symptomatic given the fluctuating nature of the disease.

We see our nasal brush-based classifier of asthma as the first step in the development of nasal biomarkers for asthma care. As with any disease, the first step is to accurately identify affected patients. The asthma gene panel described in this study provides an accurate initial path to this critical diagnostic step. With a correct diagnosis, an array of existing asthma treatment options can be considered [6]. A next phase of research will be to develop a nasal biomarker to predict endotypes and treatment response, so that asthma treatment can be targeted, and even personalized, with greater efficiency and effectiveness [51].

In summary, we demonstrated an innovative application of RNA sequencing and machine learning to identify a classifier consisting of genes expressed in nasal brushings that accurately classifies subjects with mild/moderate asthma from controls. This asthma classifier performed with accuracy across independent and external test sets, indicating reasonable generalizability across study populations and gene expression profiling modality, as well as specificity to asthma. This asthma classifier could potentially lead to the development of a nasal biomarker of asthma.

## Materials and Methods

### Study design and subjects

Subjects with mild/moderate asthma were a subset of participants of the Childhood Asthma Management Program (CAMP), a multicenter North American study of 1041 subjects with mild to moderate persistent asthma [19, 20]. Findings from the CAMP cohort have defined current practice and guidelines for asthma care and research [20]. Asthma was defined by symptoms ≥2 times per week, use of an inhaled bronchodilator ≥ twice weekly or use of daily medication for asthma, and increased airway responsiveness to methacholine (PC_20_≤12.5 mg/ml). The subset of subjects included in this study were CAMP participants who presented for a visit between July 2011 and June 2012 at Brigham and Women’s Hospital (Boston, MA), one of the eight study centers for CAMP.

Subjects with “no asthma” were recruited during the same time period by advertisement at Brigham & Women’s Hospital. Selection criteria were no personal history of asthma, no family history of asthma in first-degree relatives, and self-described Caucasian ethnicity. Participation was limited to Caucasian individuals because a concurrent independent study was planned that would compare these same subjects to 968 Caucasian CAMP subjects who participated in the CAMP Genetics Ancillary study [52]. Subjects underwent pre- and post-bronchodilator spirometry according to American Thoracic Society guidelines. Only those meeting selection criteria and with demonstrated normal lung function without bronchodilator response were considered to have “no asthma.”

### Nasal brushing and RNA sequencing

Nasal brushing was performed with a cytology brush. Brushes were immediately placed in RNALater (ThermoFisher Scientific, Waltham, MA) and then stored at 40^0^C until RNA extraction. RNA extraction was performed with Qiagen RNeasy Mini Kit (Valencia, CA). Samples were assessed for yield and quality using the 2100 Bioanalyzer (Agilent Technologies, Santa Clara, CA) and Qubit fluorometry (Thermo Fisher Scientific, Grand Island, NY).

Of the 190 subjects who underwent nasal brushing (66 with mild/moderate asthma, 124 with no asthma), a random selection of 150 subjects were *a priori* assigned as the development set (for classification model development), with the 40 remaining subjects earmarked to serve as a test set of independent subjects (for testing the classification model). To minimize potential batch effects, all samples were submitted together for RNA sequencing (RNAseq). Staff at the Mount Sinai genomics core were blinded to the assignment of samples as development or test set. The sequencing library was prepared with the standard TruSeq RNA Sample Prep Kit v2 protocol (Illumina). The mRNA libraries were sequenced on the Illumina HiSeq 2500 platform with a per-sample target of 40-50 million 100 bp paired-end reads. The data were put through Mount Sinai’s standard mapping pipeline[53] (using Bowtie [54] and TopHat [55], and assembled into gene- and transcription-level summaries using Cufflinks [56]). Mapped data were subjected to quality control with FastQC and RNA-SeQC [57]. Data were pre-processed separately for the development and test sets to avoid leakage of information across the two data sets and maintain fairness of the machine learning procedures as much as possible. Genes with fewer than 100 counts in at least half the samples were dropped to reduce the potentially adverse effects of noise. DESeq2 [23] was used to normalize the data sets using its variance stabilizing transformation method.

### VariancePartition Analysis of Potential Confounders

Given differences in age, race, and sex distributions between the asthma and “no asthma” classes, we used the variancePartition method [22] to assess the degree to which these variables influenced gene expression and potentially confounded the target phenotype (asthma status). The total variance in gene expression was partitioned into the variance attributable to age, race, and sex using a linear mixed model implemented in variancePartition v1.0.0 [22]. Age (continuous variable) was modeled as a fixed effect, while race and sex (categorical variables) were modeled as random effects. The results showed that age, race, and sex accounted for minimal contributions to total gene expression variance (**Supplementary Figure 1**). Downstream analyses were therefore performed with gene expression data unadjusted for these variables.

### Differential gene expression and pathway enrichment analysis

DESeq2 [23] was used to identify differentially expressed genes in the development set. Genes with FDR ≤ 0.05 were deemed differentially expressed, with fold change <1 implying under-expression and vice versa. To identify the functions underlying these genes, pathway enrichment analysis was performed using the Gene Set Enrichment Analysis method applied to the Molecular Signature Database (MSigDB) [24].

### Identification of the Asthma Classifier by Machine Learning Analyses of the RNAseq Development Set

To identify gene expression-based classifiers that predict asthma status, we applied a rigorous machine learning pipeline that combined feature (gene) selection [16], classification [17], and statistical analyses of classification performance [18] to the development set (**Supplementary Figure 2**). The pipeline was implemented in Python using the scikit-learn package [58]. Feature selection and classification were applied to a training set comprised of 120 randomly selected samples from the development set (n=150) as described below. For an independent evaluation of the candidate classifiers generated from the training set by this process, they were then evaluated on the remaining 30 samples (holdout set). Finally, to reduce the dependence of the finally chosen classifier on a specific training-holdout split, this process was repeated 100 times on 100 random splits of the development set into training and holdout sets. The details of the overall process as well as the individual components are as follows.

#### Feature selection

The purpose of the feature selection component was to identify subsets of the full set of genes in the development set, whose expression profiles could be used to predict the asthma status as accurately as possible. The two main computations constituting this component were (i) the optimal number of features that should be selected, and (ii) the identification of this number of genes from the full gene set. To reduce the likelihood of overfitting when conducting both these computations on the entire training set, we used a 5x5 nested (outer and inner) cross-validation (CV) setup [30] for selecting features from the training set (**Supplementary Figure 3**). The inner CV round was used to determine the optimal number of genes to be selected, and the outer CV round was used to select the set of predictive genes based on this number, thus separating the samples on which these decisions are made. The supervised Recursive Feature Elimination (RFE) algorithm [59] was executed on the inner CV training split to determine the optimal number of features. The use of RFE within this setting enabled us to identify groups of features that are collectively, but not necessarily individually, predictive. Specifically, we used the L2-regularized Logistic Regression (LR or Logistic) [60] and SVM-Linear (kernel) [61] classification algorithms in conjunction with RFE (combinations henceforth referred to as LR-RFE and SVM-RFE respectively). For this, for a given inner CV training split, all the features (genes) were ranked using the absolute values of the weights assigned to them by an inner classification model, trained using the LR or SVM algorithm, over this split. Next, for each of the conjunctions, the set of top-k ranked features, with k starting with 11587 (all filtered genes) and being reduced by 10% in each iteration until k=1, was considered. The discriminative strength of feature sets consisting of the top k features as per this ranking was assessed by evaluating the performance of the LR or SVM classifier based on them over all the inner CV training-test splits. The optimal number of features to be selected was determined as the value of k that produces the best performance. Next, a ranking of features was derived from the outer CV training split using exactly the same procedure as applied to the inner CV training split. The optimal number of features determined above was selected from the top of this ranking to determine the optimal set of predictive features for this outer CV training split. Executing this process over all the five outer CV training splits created from the development set identified five such sets. Finally, the set of features (genes) that was common to all these sets (i.e. in their intersection/overlap), which is expected to yield a more robust feature set than the individual outer CV splits, was selected as the predictive gene set for this training set. One such set was identified for each application of LR-RFE and SVM-RFE to the training set.

#### Classification analyses

Once predictive gene sets had been selected from feature selection, four global classification algorithms (L2-regularized Logistic Regression (LR or Logistic) [60], SVM-Linear [61], AdaBoost [62], and Random Forest (RF) [63]) were used to learn *intermediate classification models* over the training set. These intermediate models were then applied to the corresponding holdout set to generate probabilistic asthma predictions for the samples. An optimal threshold for converting these probabilistic predictions into binary ones (higher than threshold=asthma, lower than threshold=no asthma) was then computed as the threshold that yielded the highest classification performance on the holdout set. This optimization resulted in the *proposed classification models*.

#### Statistical analyses of classification performance

After the above components were run on 100 training-holdout splits of the development set, we obtained 100 proposed classification models for each of eight feature selection-global classification combinations (two feature selection algorithms (LR-RFE and SVM-RFE) and four global classification algorithms Logistic, SVM-Linear, AdaBoost and RF). The next step of our pipeline was to determine the best performing combination. Instead of making this determination just based on the highest evaluation score, as is typically done in machine learning studies, we utilized this large population of models and their optimized holdout evaluation scores to conduct a statistical comparison to make this determination. Specifically, we applied the Friedman test followed by the Nemenyi test [18, 64] to this population of modules and their evaluation scores. These tests, which account for multiple hypothesis testing, assessed the statistical significance of the relative difference of performance of the combinations in terms of their relative ranks across the 100 splits.

#### Optimization for parsimony

For a phenotype classifier, it is essential to consider parsimony in model selection (i.e. minimize number of features (i.e. genes)) to enhance its clinical utility and acceptability. To enforce this for our asthma classifier, an adapted performance measure, defined as the absolute performance measure (F-measure) divided by the number of genes in that model, was used for the above statistical comparison, i.e. as input to the Friedman-Nemenyi tests. In terms of this measure, a model that does not obtain the best performance measure among all models, but uses much fewer genes than the others, may be judged to be the best model. The result of the statistical comparison using this adapted measure was visualized as a Critical Difference plot [18] (**Supplementary Figure 4**), and enabled us to identify the best combination of feature selection and classification method as the left-most entry in this plot.

#### Final model development

The final step in our pipeline was to determine the representative model out of the 100 learned the above best combination by finding which of these models yielded the highest evaluation measure (F-measure; **Box 1** and **Supplementary Figure 5**). In case of ties among multiple candidates, the gene set that produced the best average asthma classification F-measure across all four global classification algorithms was chosen as the gene set constituting the representative model for that combination. This analysis yielded the representative gene set, global classification algorithm, and the optimized asthma classification threshold. The asthma classifier was built by training the global classification algorithm to the expression profiles of the representative gene set, and using the optimized threshold for classifying samples positive/negative for asthma.

### Evaluation of the Asthma Classifier in an RNAseq test set of independent subjects

The asthma classifier identified by our machine learning pipeline was then tested on the RNAseq test set (n=40) to assess its performance in independent subjects. F-measure was used as the primary measure for classification performance, as described in **Box 1** and **Supplementary Figure 5**. AUC, PPV and NPV were additionally calculated for context.

### Comparison of Performance to Alternative Classification Models

Although our classifier was identified using a rigorous machine learning methodology, the pipeline explored several other models from all combinations of feature selection and global classification methods. Thus, we compared the performance of our classifier with all these other possible classifiers.

Also, our methodology was not the only way to develop gene expression-based classifiers. Thus, we also compared the classifier’s performance with several other valid methods by applying our machine learning pipeline with the feature (gene) selection step replaced with the following alternatively determined gene sets: (1) all filtered RNAseq genes, (2) all differentially expressed genes, and (3) known asthma genes from a recent review of asthma genetics [33]. To maintain consistency with the machine learning pipeline-derived models, each of these gene sets was run through the same pipeline (**Supplementary Figure 2** with the feature selection component turned off) to identify the best performing global classification algorithm and the optimal asthma classification threshold for this predetermined set of features. The algorithm and threshold were used to train each of these gene sets’ representative classification model over the entire development set, and the resulting models for each of these gene sets were then evaluated on the RNAseq test set. Finally, as a baseline representative of alternative sparse classification algorithms, which represent a one-step option for doing feature selection and classification simultaneously, we also trained an L1-regularized logistic regression model (L1-Logistic) [34] on the development set and evaluated it on the RNAseq test set.

### Comparison of Performance to Permutation-based Random Models

To determine the extent to which the performance of all the above classification models could have been due to chance, we compared their performance with that of their random counterpart models (**Supplementary Figures 6 and 7**). These counterparts were obtained by randomly permuting the labels of the samples in the development set and executing each of the above model training procedures on these randomized data sets in the same way as for the real development set. These random models were then applied to each of the test sets considered in our study, and their performances were also evaluated in terms of the same measures. For each of real models tested in our study, 100 corresponding random models were learned and evaluated as above, and the performance of the real models was compared with the average performance of the corresponding random models.

### Evaluation of the asthma classifier in external independent asthma cohorts

To assess the generalizability of the asthma classifier to other populations, microarray-profiled data sets of nasal gene expression from two external asthma cohorts-- Asthma1 (GSE19187) [35] and Asthma2 (GSE46171) [36] (**Supplementary Table 3**)-- were obtained from NCBI Gene Expression Omnibus (GEO) [65]. For each of these data sets, we obtained their probe-level normalized versions from GEO, and then obtained gene-level expression profiles by averaging the normalized expression of all the probes corresponding to the same gene. The probe-to-gene mappings were obtained from the microarray platform (GPL) files also available from GEO. The asthma classifier was then applied to these gene-level data sets and its performance evaluated on these external asthma cohorts.

### Evaluation of the asthma classifier in external cohorts with other respiratory conditions

To assess the classifier’s ability to distinguish asthma from respiratory conditions that can have overlapping symptoms with asthma, i.e. its specificity to asthma, microarray-profiled data sets of nasal gene expression were also obtained for five external cohorts with allergic rhinitis (GSE43523) [39], upper respiratory infection (GSE46171) [36], cystic fibrosis (GSE40445) [40], and smoking (GSE8987) [12] (**Supplementary Table 4**). Gene-level expression data sets were obtained for these cohorts using the same methodology as described above. The asthma classifier was then applied to these data sets, and its performance evaluated on these external cohorts with non-asthma respiratory conditions.

### Data availability

Data and code for this study (doi:10.7303/syn9878922) will be made publically available upon publication via Synapse, a software platform for open, reproducible data-driven science, at https://www.synapse.org/#!Synapse:syn9878922/files/

## Declarations

### Ethics approval and consent to participate

The institutional review boards of Brigham & Women’s Hospital and the Icahn School of Medicine at Mount Sinai approved the study protocols. Written informed consent was obtained from all subjects and all research was performed in accordance with relevant guidelines and regulations.

### Author contributions

SB directed the study. SB and AJR directed the recruitment of subjects and sample collection. BAR and STW provided guidance for access to subjects. EES advised on sequencing strategy. SB curated the clinical data. SB, GP, EES, OPP and MEA designed and performed the statistical and computational analyses. SB and GP wrote the manuscript. SB, GP, OPP, AJR, MEA, GEH, BAR, STW, and EES edited the manuscript. All authors contributed significantly to the work presented in this paper.

## Acknowledgments

We thank Yoojin Chun, Kathryn Paul, Laura Ting, Anne Plunkett, Nancy Madden, Ann Fuhlbrigge, Kelan Tantisira, Dan Cossette, Aimee Garciano, and Roxanne Kelly for their assistance and support with recruitment, specimen collection, and sample processing. We thank Robert Griffin and Ana Stanescu for critically reviewing the paper.

## Supplementary Materials

**Supplementary Figure 1:** variancePartition analysis of the RNAseq development set.

**Supplementary Figure 2:** Visual description of the machine learning pipeline used to select predictive features (genes) and develop classification models based on them in the RNAseq development set.

**Supplementary Figure 3:** Visual description of the feature (gene) selection component of the machine learning pipeline.

**Supplementary Figure 4:** Critical Difference plots demonstrating results of the statistical comparison of the performance of 100 asthma classification models obtained by various combinations of feature selection and global classification algorithms in terms of the classification performance and parsimony (numbers of genes included) of the models.

**Supplementary Figure 5:** Evaluation measures for classification models[31].

**Supplementary Figure 6:** Performance of permutation-based random classification models in test sets of independent subjects with asthma and controls.

**Supplementary Figure 7:** Performance of permutation-based random classification models in test sets of independent subjects with non-asthma respiratory conditions and controls.

**Supplementary Figure 8:** Distribution of DESeq2 FDR values of differential expression in the asthma classifier (blue bars) vs. other genes in the RNAseq development set (coral bars).

**Supplementary Table 1:** Lists of over- and under-expressed genes and pathways in asthma cases compared to controls (in different tabs of this file). Differentially expressed genes were identified using DESeq2 [23] applied to the development set, and enriched pathways were identified from the Molecular Signature Database [24], both using an upper FDR threshold of 0.05.

**Supplementary Table 2:** List of known asthma-associated genes from a review of asthma genetics [33] that overlap with genes in our RNAseq data sets.

**Supplementary Table 3:** Characteristics of the external asthma cohorts used in the evaluation of the asthma classifier.

**Supplementary Table 4:** Characteristics of the external cohorts with non-asthma respiratory conditions and controls used in the evaluation of the asthma classifier.

**Supplementary Table 5:** Basic functional annotations and references for asthma classifier genes that have been studied in the context of asthma and airway inflammation

